# Dramatic evolution of body length due to post-embryonic changes in cell size in a newly discovered close relative of *C. elegans*

**DOI:** 10.1101/181107

**Authors:** Gavin C. Woodruff, Patrick C. Phillips

## Abstract

Understanding morphological diversity—and morphological constraint—has been a central question in evolutionary biology since its inception. Nematodes of the genus *Caenorhabditis*, which contains the well-studied model system *C. elegans*, display remarkable morphological consistency in the face of extensive genetic divergence. Here, we provide a description of the broad developmental patterns of a recently discovered species, *C.* sp. 34, which was isolated from fresh figs in Okinawa and which is among the closest known relatives of *C. elegans*. *C.* sp. 34 displays an extremely large body size and can grow to be nearly twice as long as *C. elegans* and all other known members of the genus. Observations of the timing of developmental milestones reveal that *C.* sp. 34 develops about twice as slowly as *C. elegans*. Measurements of embryo and larval size show that the size difference between *C.* sp. 34 and *C. elegans* is largely due to post-embryonic events, particularly during the transition from larval to adult stages. This difference in size is not attributable to differences in germ line chromosome number or the number of somatic cells. The overall difference in body size is therefore largely attributable to changes in cell size via increased cytoplasmic volume. Because of its close relationship to *C. elegans*, the distinctness of *C.* sp. 34 provides an ideal system for the detailed analysis of evolutionary diversification. The context of over forty years of *C. elegans* developmental genetics also reveals clues into how natural selection and developmental constraint act jointly to promote patterns of morphological stasis and divergence in this group.

---

It is natural for evolutionary biologists to focus on change; the more dramatic, the better. However, we expect species to accumulate substantial differences from one another over time even in the absence of natural selection[1]. In fact, even across fairly diverse groups, the predominant pattern of evolution is one of constrained variation in morphological diversity rather than diversification per se[2]. For the last 40 years, the biological bases of limits to macroevolutionary variation have been hotly debated[3-6]. In the early phases of this discussion, evolutionary geneticists tended to argue that long term limits to variation must be generated by stabilizing selection in which the natural tendency for species to move apart from one another in morphological space due to the accumulation of new mutations via genetic drift is strongly counterbalanced by natural selection against individuals that do not adhere to an optimal phenotype[4]. In contrast, evolutionary developmental biologists and paleontologists often argued that development systems themselves constrain the actual production of variation that is the basis of evolutionary change, such that species that share common developmental regulatory systems would be expected to show limited phenotypic difference from one another[5]. In the intervening years, it has become clear that the actual diversity that we observe in nature must somehow be a balance between these different sources of constraint[2, 6].

Nematodes are a particularly compelling example of extremely constrained morphological evolution. For instance, species within the genus *Caenorhabditis*, which includes the important *C. elegans* model system, display such little morphological diversity that they are essentially impossible to tell apart except in a few finer details of male tail morphology[7, 8]. In general, species can only be defined via their ability to cross with one another[8]. Yet this morphological conservatism is in stark contrast to amount of diversity observed at the level of DNA sequence.

Here, different species within this group are as divergent from one another as mice are from humans[9]. Nematodes are famous for having a very stereotypical pattern of development, with a largely fixed lineage of cell division events and number of adult somatic cells[10, 11]. Is the constrained pattern of morphological diversity observed within this genus generated by development or selection? Here, we test this hypothesis using the developmental characteristics of a recently discovered relative of *C. elegans*, *C.* sp. 34 (whose discovery, taxonomic status, and genome sequence have been recently formally described[12]). In addition to exhibiting exceptional differences in body size and other morphological characteristics, this species is also distinctive from other *Caenorhabditis* species in its developmental rate and ecological niche[12]. We describe a number of these features, with a particular focus on examining the proximal causes of the extreme difference in body size.

## Results

### A morphologically novel species of fig-associated nematode is in *Caenorhabditis*

C. sp. 34 was originally isolated from the fresh figs of Ficus septica in Okinawa, Japan, and phylogenetic analyses place this species among the closest reported relatives of *C*. elegans[12] (Fig. 1; Supplemental Document 1). Surprisingly, *C*. sp. 34 is an exceptional Caenorhabditis species in a number of respects. In contrast to other morphologically indistinguishable species of the elegans group, they are huge in size, on average 64% longer than its close relative *C.* elegans[12] (Fig 1A; Fig. 5, see below). *C.* sp. 34 females have a distinctive, stumpy tail morphology, with a much shorter tail spike than those of *C. elegans* hermaphrodites[12] (Fig. 2a-c). In addition, *C.* sp. 34 has enormous sperm that are on average three times longer in diameter than those of *C. elegans* (Fig. 2d-f). *C.* sp. 34 also develops very slowly, with a generation time nearly twice as long as *C. elegans* (Fig. 3, see below). Furthermore, mating tests between *C. elegans* and *C.* sp. 34 yielded no viable progeny[12] (Supplemental Document 2). *C.* sp. 34 is also exceptional in its ecological niche, with proliferating animals being found in fresh figs[12] (Fig. 1b), whereas most *Caenorhabditis* animals are associated with rotting plant material[7].

**Figure 1.**
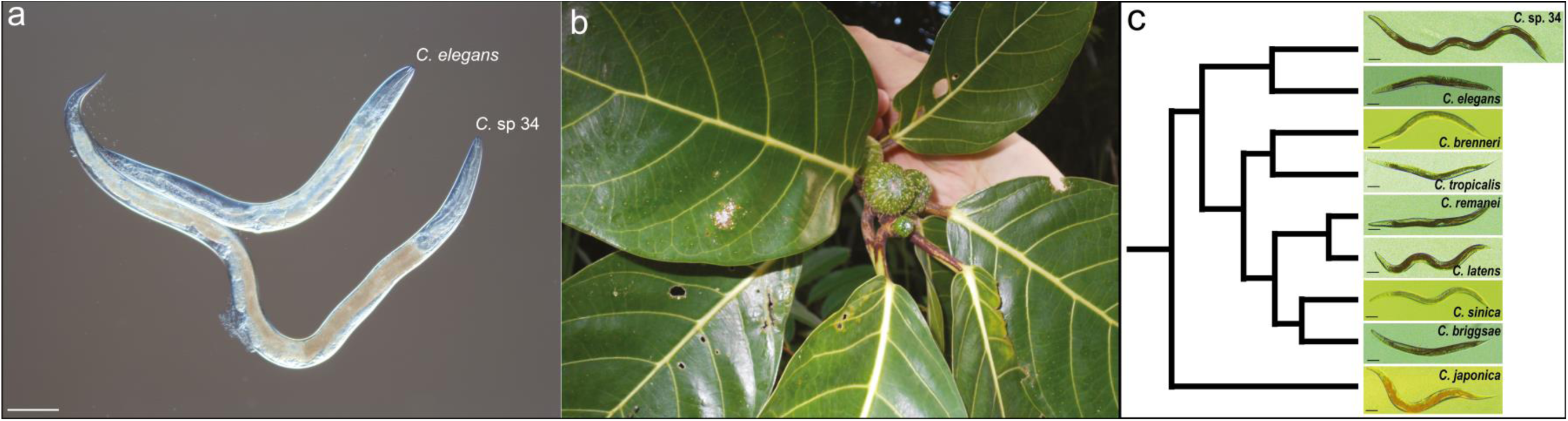
*C.* sp. 34 is a morphologically and ecologically distinct species of *Caenorhabditis*. (a) *C.* sp. 34 is longer than *C. elegans*. (b) *C.* sp. 34 is associated with the fresh figs of *Ficus septica*, in contrast to most *Caenorhabditis* species, which are associated with rotting plant material[7]. (c) *C.* sp. 34 is a morphologically exceptional *Caenorhabditis*. Age-synchronized *Caenorhabditis* females/hermaphrodites across nine species shows *C.* sp. 34 to be highly derived in its body length. The cladogram follows the analysis in[12]. *C. latens* has been previously shown to be the sister species of *C. remanei*[8, 13]. All scale bars are 100 microns.

**Figure 2.**
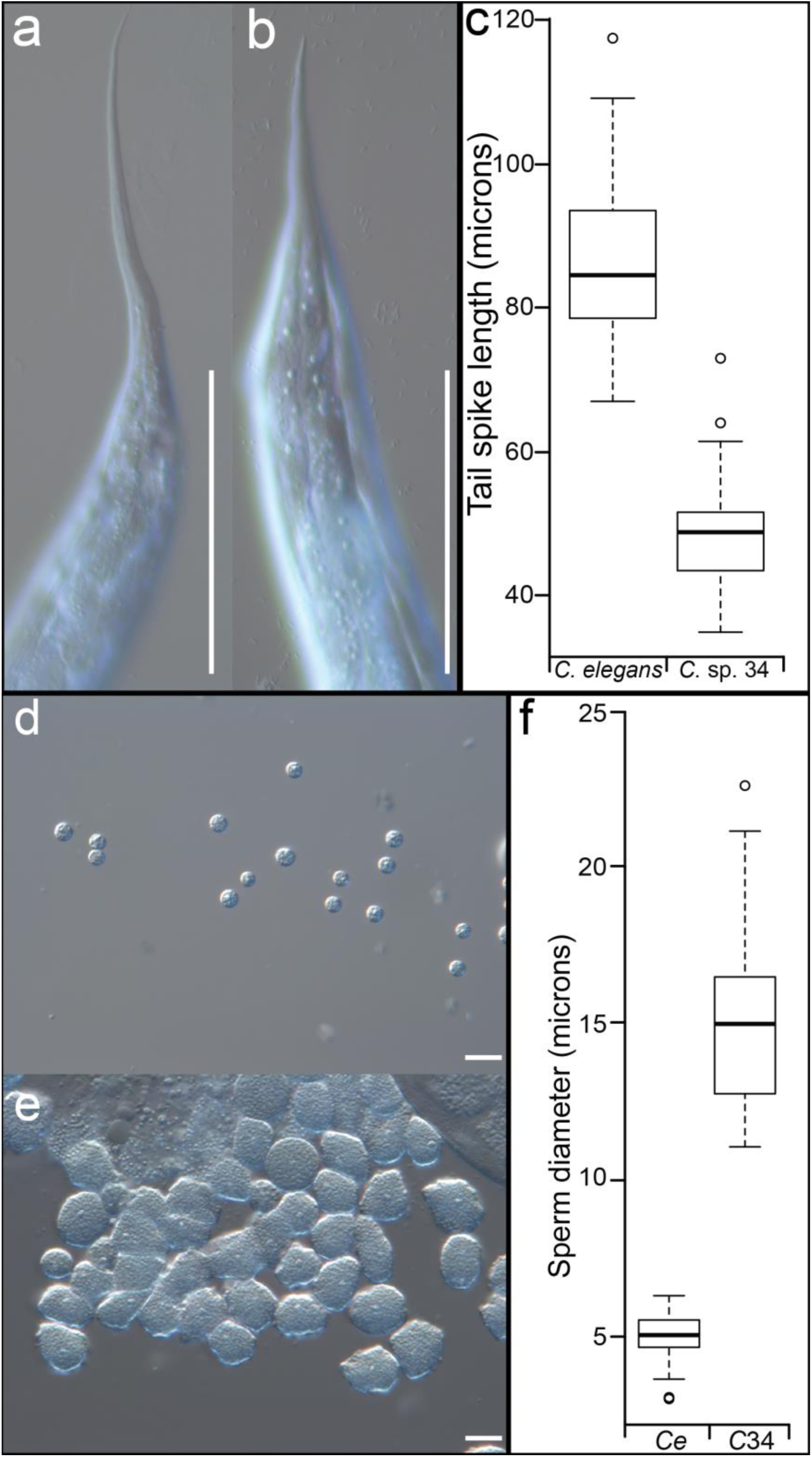
*C.* sp. 34 has small female tail spikes and giant sperm. *C*. *elegans* N2 hermaphrodite (a) and *C.* sp. 34 NKZ1 female (b) tail spikes. *C*. *elegans* hermaphrodite tail spikes had an average length of 86.5 microns (N=43 worms, **±** 5.3 SDM), whereas *C.* sp. 34 female tail spikes had an average length of 48.3 microns (Mann-Whitney U p<0.0001, N=41 worms, **±** 3.7 SDM). Scale bars are 100 microns in both photos. (c) Quantification of tail spike length. *C*. *elegans* n=43, *C.* sp. 34 n=41. (d) Sperm dissected from *C*. *elegans (fog-2)* males. (e) Sperm dissected from *C.* sp. 34 NKZ1 males. (f) Quantification of sperm size diameter. *C*. *elegans (fog-2)* male sperm had an average diameter of 5.06 microns (N=56 sperm, **±** 0.36 SDM), whereas *C.* sp. 34 sperm had an average diameter of 15.07 microns (Mann-Whitney U p<0.0001, N=45 sperm, **±** 1.37 SDM). Scale bars are 10 microns in both photos.

**Figure 3.**
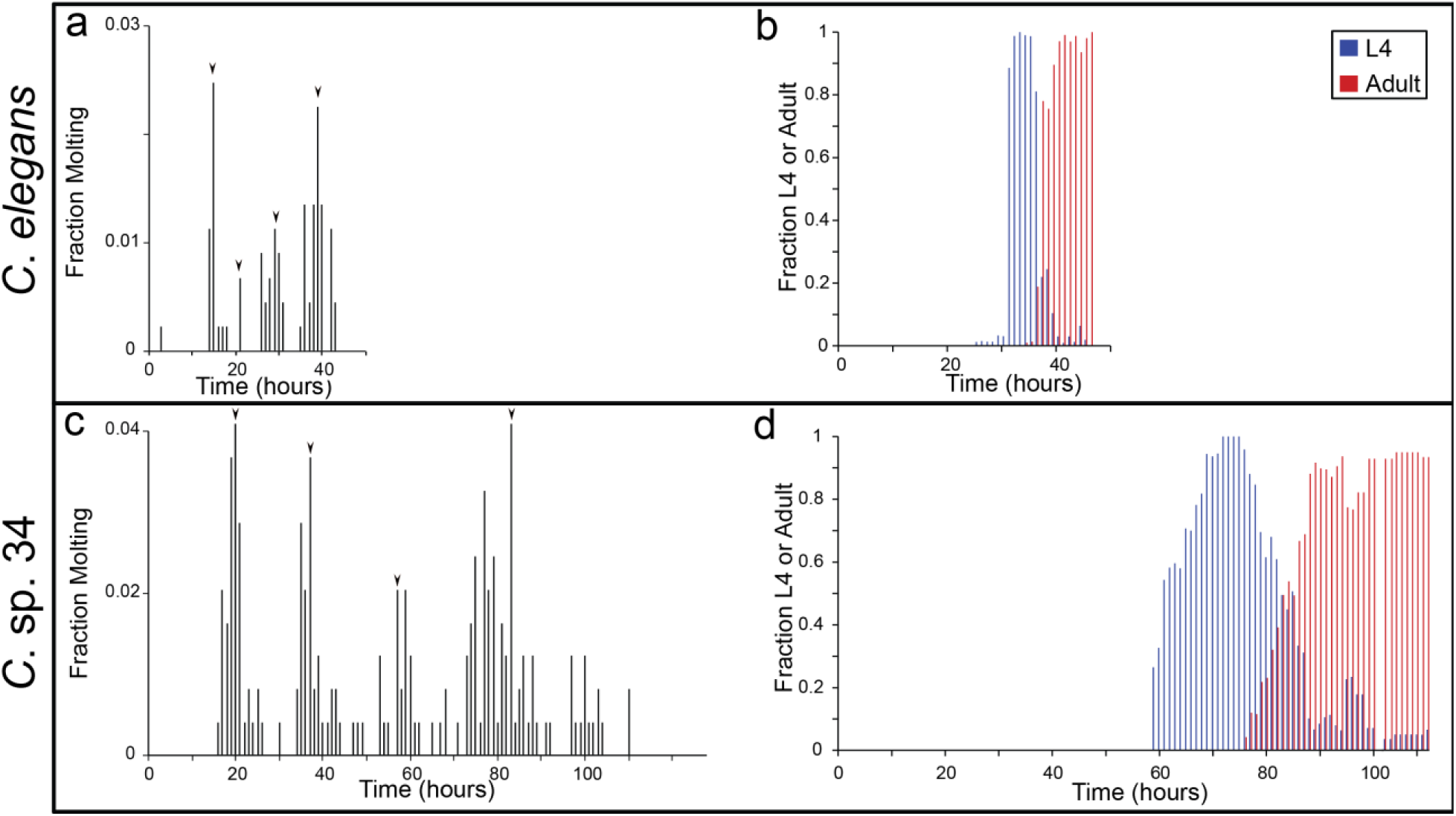
*C.* sp. 34 develops more slowly than *C. elegans*. Synchronized populations of *C. elegans* (a, b) and *C.* sp. 34 (c, d; for both species, average N worms=107 **±**32.2 SDM; range= 23-445), were monitored hourly for the fraction of actively molting animals (a, c) and the fraction of L4 larvae and adults (b, d). Populations were synchronized at the L1 larval stage twelve hours apart and monitored concurrently to capture the full progression of development. Arrowheads in (a) and (c) represent the maximal molting fractions corresponding to the likely major molting events.

### *C.* sp. 34 develops slowly

*C. elegans* typically takes about two days to develop at 25°C[14]. However, it was readily apparent that *C.* sp. 34 has a much slower developmental rate. This was quantified by examining the fraction of animals actively molting and the number of animals in the L4 and adult stages (which can be easily ascertained morphologically) over time (Figure 3).The four larval molts are highly conserved across nematodes[15], and this is reflected in the periodicity of the molting fraction of both *C. elegans* and *C.* sp. 34 (Figure 3a, 3c). *C. elegans* had maximal molting fractions at 15, 21, 28 and 39 hours past L1 synchronization (Figure 3a). Conversely, *C.* sp. 34 had maximal molting fractions at 21, 38, 58, and 76 hours (Figure 3c), revealing a developmental rate that is about twice as slow. This difference is also apparent in the proportion of L4 - and adult-like animals over time. The maximal L4 and adult fractions occur in *C. elegans* at 34 and 46 hours past L1 synchronization, whereas in *C.* sp. 34 they are at 72 and 106 hours (Figure 3b, 3d). In addition, there is much more variation in developmental rate in *C.* sp. 34 than *C. elegans*. The amount of time in which L4 larvae were observed was over twice as long in *C.* sp. 34 (53 hours) than in *C. elegans* (21 hours).

### *C.* sp. 34 is not polyploid

One explanation then for the increased size of *C.* sp. 34 is of a chromosome or genome duplication event. For instance, polyploid strains of *C. elegans* were initially generated to show that the X chromosome:autosome ratio was the major determiner of sex[16, 17], but it was also noted that polyploid animals are larger than wild-type[17, 18]. Ploidy can be easily ascertained by examining DNA-stained oocytes, which in *Caernorhabditis* arrest in prophase I prior to maturation[19], allowing for chromosomes to be easily visualized (Figure 4). In all *C.* sp. 34 specimens where the chromosomes in diakinesis-stage oocytes were apparent (n=29), six chromosomes were observed. This was also true of *C*. *elegans* (n=15). This result consistent with recently reported genomic and microscopic data[12]. In many *C.* sp. 34 animals, germ line and oocyte nuclear abnormalities were observed (Supplemental Figure 1). This may reflect oocyte endoreplication or chromosome condensation in the unmated animals used for microscopy, which can also be observed in older, sperm-depleted *C*. *elegans* hermaphrodites[20]. This may also reflect nutritional deficiencies in standard *C. elegans* laboratory confidtions for *C.* sp. 34, as starvation conditions are known to affect the germline in *C*. *elegans*[21].

**Figure 4.**
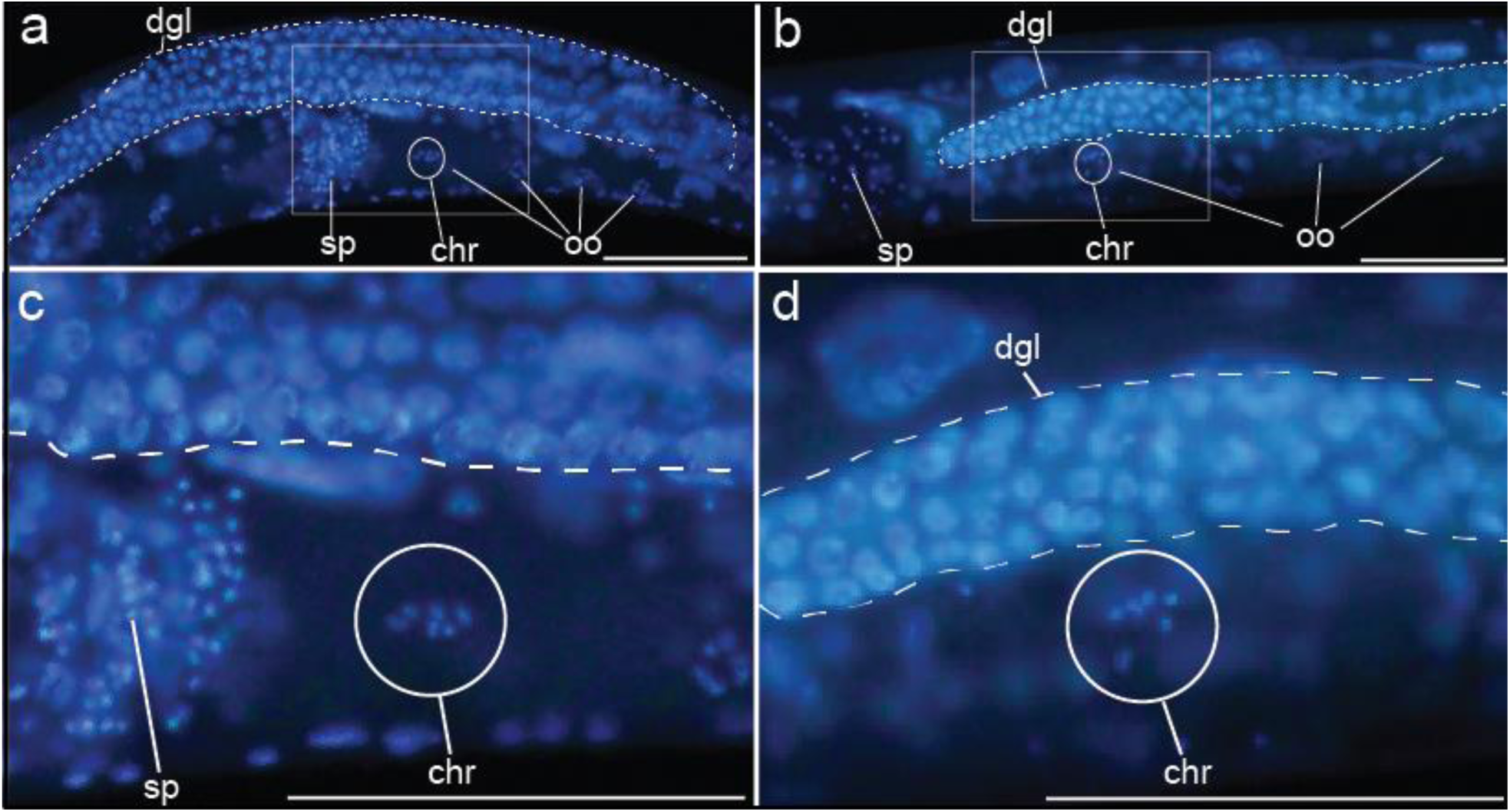
*C.* sp. 34 and *C. elegans* have the same number of chromosomes. DNA stained *C. elegans* (A, C) and *C.* sp. 34 (B, D) reveal that late-prophase I oocytes contain six chromosomes (chr, encircled). Also of note is the reduced *C.* sp. 34 gonad relative to *C. elegans*. Scale bars represent 100 microns in all photos. dgl, distal germ line. sp, sperm. oo, oocyte.

### *C.* sp. 34 length difference is largely due to post-embryonic events

To investigate the developmental basis of the size difference between *C.* sp. 34 and *C*. *elegans*, length was measured over time and developmental stage (Figure 5). Despite being 64% longer on average than *C*. *elegans* at four days after egg-laying, *C.* sp. 34 embryos are are only 19% longer than *C. elegans* embryos (Fig. 5a-c). Thus, it appears that a substantial portion of the length difference between these species is due to post-embryonic events. However, as development is delayed in *C.* sp. 34 compared to *C*. *elegans* (Fig. 3), length comparisons at the same time are problematic. To address this, the lengths of animals were compared at similar developmental stages (Fig. 5d-j). Although *C.* sp. 34 is longer than *C*. *elegans* at all developmental stages (Fig. 5j), between the L3 and adult stages the average length difference grows from 33% to 64%. Thus, much of the difference in length between species is developmentally regulated during the larva-to-adult transition. In addition, although *C.* sp. 34 adults are observed to be significantly wider than *C. elegans* (Mann-Whitney U p=0.02), they are nominally wider on average by only four microns (Fig. 5k). The size difference between these species is then dominated by length.

**Figure 5.**
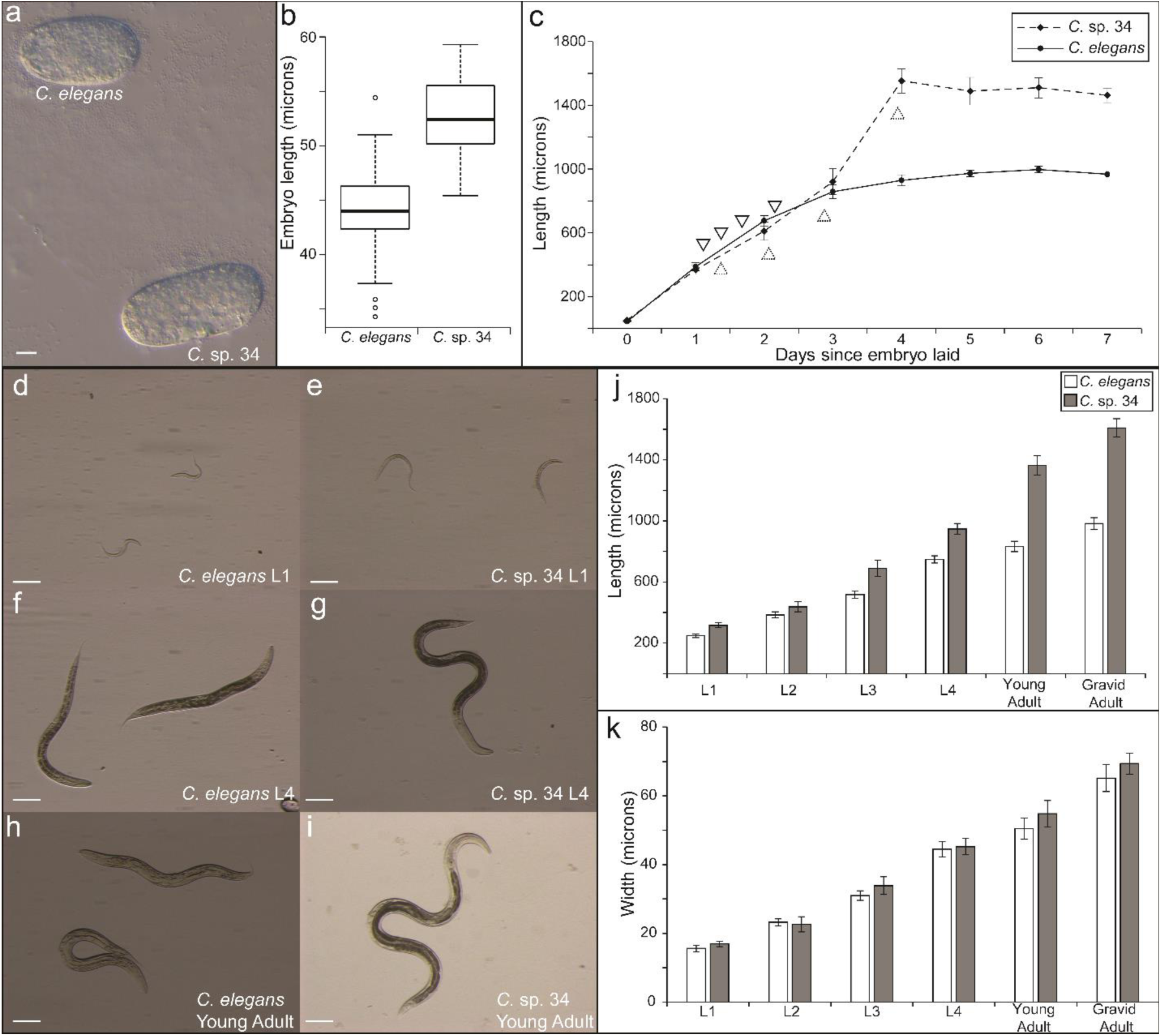
The length difference between *C. elegans* and *C.* sp. 34 is largely due to post-embryonic events. (a) *C. elegans* and *C.* sp. 34 embryos. Scale bar=10 microns. (b) Boxplot comparing embryo length (n=61 for *C*. *elegans*; n=35 for *C.* sp. 34; Mann-Whitney U p<0.0001). *C.* sp. 34 embryos are on average 19% longer than *C*. *elegans* embryos. (c) Comparison of body length size over time in populations of *C.* sp. 34 and *C. elegans* synchronized as embryos (average N worms=21 **±**3.7 SDM; range=11-36). Data at time “0” is the same as in (b). Arrows correspond to estimates of larval molts in *C. elegans* (pointing down) and *C.* sp. 34 (pointing up) as determined in a Fig. 3. (d-i) Images of *C*. *elegans* (d, f, h) d *C.* sp. 34 (e, g, i) at developmentally comparable stages. Scale bars correspond to 100 microns in all panels. (j) Comparison of body length at developmental stages. *C.* sp. 34 is significantly longer than *C*. *elegans* at all stages (Mann-Whitney U p <0.003 for all stages), but a 27% length difference at the L1 stage grows to a 64% difference in adults (Average N worms=33 **±**4.1 SDM; range=16-41). (k) Comparison of body width of same animals as in (j). The width of *C.* sp. 34 is comparable to *C*. *elegans* at all developmental stages. Error bars represent one standard deviation of the mean in panels (c, j-k).

### *C.* sp. 34 size difference is due to differences in cell size and not cell number

All differences in body size must be due to differences in cell number, cell size, or both. To distinguish between these possibilities, somatic nucleus numbers (used as a proxy for cell number) were hand-counted in unmated *C.* sp. 34 adult females and *C. elegans* (*fog-2*) adult pseudo-females. The *fog-2* mutation was used in order to provide comparable specimens that lacked self-embryos, which had the potential to contribute to error in somatic nucleus number estimation. In addition, *fog-2* mutants have no known somatic defects[22]. Germ cells were not counted as *C.* sp. 34 germ lines are reduced relative to *C*. *elegans* (Fig. 4, Supplemental Figure 1), and it is unlikely that this tissue would contribute to the length difference. No significant difference in somatic nuclei number between *C*. *elegans* and *C.* sp. 34 was observed (Mann-Whitney U p =0.09; Fig. 6a). Although *C.* sp. 34 showed a nominal (yet not significant) difference in average nucleus number (23 more nuclei than *C. elegans* on average), this is not enough to explain a >60% difference in adult length, as the *C. elegans* hermaphrodite has about 959 somatic nuclei (which is largely invariant;[14]). Thus it is likely that differences in cell size, and not cell number, mostly explain the difference in length between *C.* sp. 34 and *C. elegans*.

**Figure 6.**
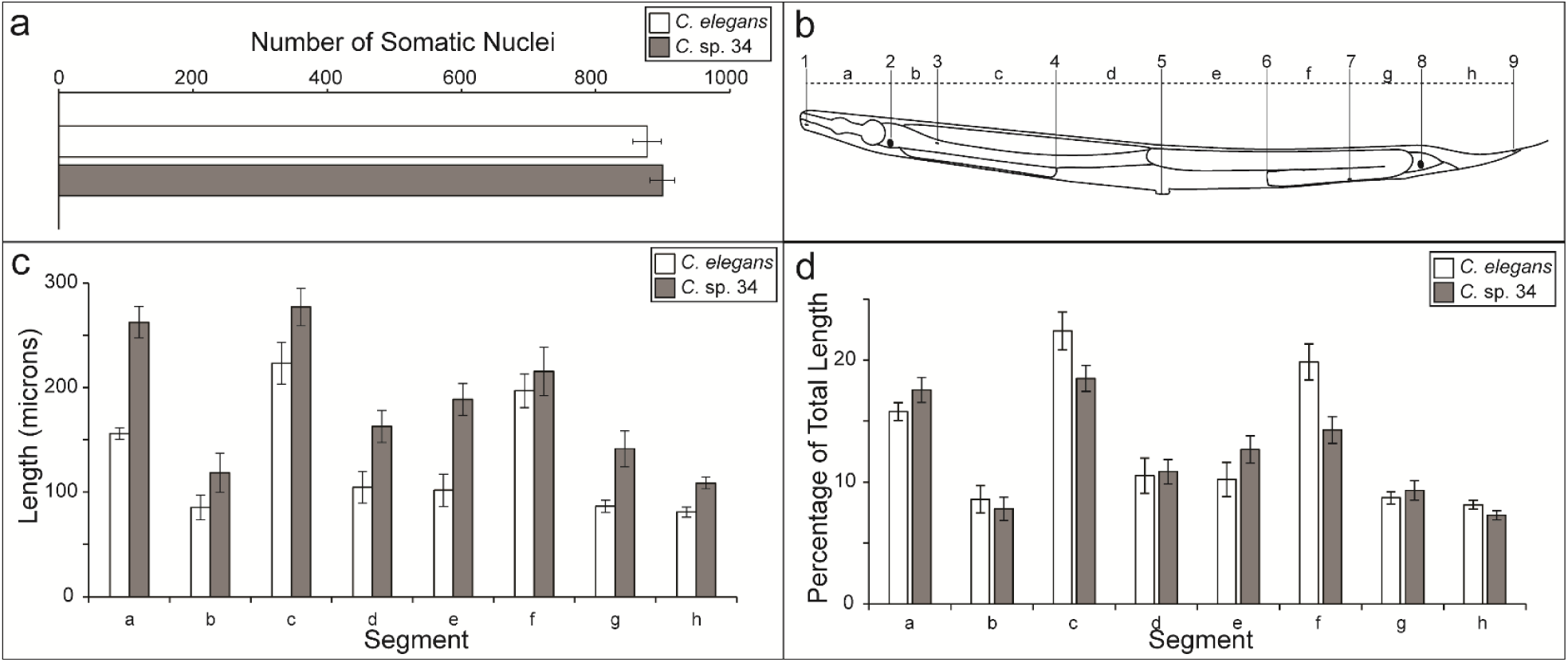
The size difference between *C. elegans* and *C.* sp. 34 is largely due to differences in cell size and not cell number. (a) Total number of somatic nuclei in young adult unmated *C.* sp. 34 and *C*. *elegans* (*fog-2)* females. *fog-2* animals make no sperm nor self-progeny but are somatically identical to wild-type hermaphrodites. No significant difference between nucleus number was observed (Mann-Whitney U p=0.09). n=13 for both species. (b) Schematic of morphological markers used to measure length segments (adapted from WormAtlas[23]). The distances (letters) between homologous nuclei (numbers) were compared between young adult *C.* sp. 34 and *C*. *elegans* (*fog-2)* females. The specific morphological landmarks used are detailed in the experimental procedures. (c) The length between homologous morphological markers in *C*. *elegans* and *C.* sp. 34. All homologous segments are significantly longer in *C.* sp. 34, with the exception of segment f. n= 15 for *C. elegans*, n=16 for *C.* sp. 34. (d) The percentage of the total body length of given homologous segments in *C.* sp. 34 and *C*. *elegans*. Same data as in (c). Although largely comparable in proportion, segments a, c, e, f, and h consist of a significantly different percentage of the total body size between *C*. *elegans* and *C.* sp. 34 (Mann-Whitney U p-values for segments a-h: 0.009 (segment a); 0.29 (segment b); 0.00031 (segment c); 0.89 (segment d); 0.0063 (segment e); <0.0001 (segment f); 0.36 (segment g); 0.0036 (segment h). Error bars represent one standard deviation of the mean in panels (a, c-d).

To quantify differences in cell size, the distances between homologous morphological landmarks in *C.* sp. 34 and *C. elegans* (*fog-2*) adult females were measured (Fig 6b-d). If the number of cells between these species is comparable, and their total length is different, then there should be differences in the distances between homologous landmarks. Furthermore, it is possible that the length difference between species is dominated by the size increase of a particular organ (such as the pharynx). If this is true, then the proportion of the total body size represented by the difference between two homologous markers should be different between species.

Indeed, the distances between homologous markers are greater in *C.* sp. 34 than *C. elegans* for every pair examined except for one (Fig. 6c). This is consistent with *C.* sp. 34 having larger cells than *C. elegans*. The one pair of morphological markers that are similarly spaced apart in *C*. *elegans* and *C.* sp. 34 is the posterior spermatheca and a posterior ventral cord neuronal nucleus (VD11; Fig. 6c). This similarity in length could be due to differences in gonad morphology, as this is influenced by the germline, which is often reduced in size or otherwise defective in *C.* sp. 34 (Fig. 4; Supplemental Figure 1). In addition, the proportion of the total body size represented by the distance between two given homologous markers is largely comparable between species (Fig. 6d). This suggests that there is a global increase in cell size in *C.* sp. 34 compared to *C. elegans*.

## Discussion

Genetic diversity drives phenotypic change. More than a hundred years of investigation has demonstrated that a multitude of quantitative traits can be readily transformed under natural and artificial selection[24-26]. Even substitutions of one or two simple genetic elements have been found to promote profound phenotypic changes within species[27]. Thus, we would expect that a high degree of genetic diversity should provide ample material for the evolution of morphological diversity. The persistence of morphological stasis across long periods of time therefore remains an apparent paradox in evolutionary biology[2, 28]. Although often framed with respect to the fossil record, this observation also holds in extant taxa. Since the onset of the molecular era, the pace of descriptions of cryptic species has been exponential[29], and the frequency of such species is not limited by phylum or geographical region[30]. Morphological stasis in the face of genetic divergence is thus likely quite common and remains a largely ignored problem in evolutionary biology.

Such stasis is often explained by long-term stabilizing selection, which purges divergent unfit forms from populations and reduces phenotypic variation[4, 31]. Stabilizing selection is thought to be acting in most populations mainly because most organisms appear to be well-adapted to their environments and thus some form of stabilizing selection must be ongoing[32]. However, others have argued that selection alone cannot explain the paradox of stasis[2, 33]. One alternative explanation often invoked is the notion of developmental constraint[5]. Here, phenotypic variation is limited by biases in the structure of the developmental genetic system itself, and divergence fails to occur because certain classes of phenotypes are not accessible to selection. This explanation is appealing due to the multitude of established such biases in developmental trajectories[34]; multiple examples of convergent phenotypic evolution promoted by the same nucleotide substitution, suggestive of limitations to the number of paths evolution can take[27, 35]; and the prevalence of correlated traits, consistent with genetic constraints that influence the range of possible phenotypes[2, 6]. Still others have suggested that the observation of long-term stasis can be resolved by invoking an incomplete fossil record and the rapid turnover of locally adapted forms[6, 28, 33, 36], as well as the difficulty of empirically detecting acting stabilizing selection when populations are close to trait optima[37]. Indeed, it is likely that a plurality of causes, including the joint action of selection and developmental constraint, contribute to patterns of long-term morphological stasis[2, 6].

The nematode genus *Caenorhabditis* represents a striking example of phenotypic constancy in the face of genetic change. Despite about twenty million years of evolution[38], the twelve reported species of the Elegans group of *Caenorhabditis* are morphologically indistinguishable (Fig. 1C), and mating tests must be used to delineate them from one another[7, 8, 39]. This phenotypic constancy persists within the context of extreme genetic divergence within and between species. The hermaphroditic *C. elegans* and *C. briggsae* share about the same degree of genetic divergence as human and mouse[9]. The male/female species *C. brenneri* and *C. remanei* harbor tremendous intraspecies polymorphism and are among the most diverse metazoans known[40], despite their cryptic species status[41]. Furthermore, this morphological constancy has persisted despite ecological diversification in this group. Many *Caenorhabditis* species are generalists that are globally distributed and are found associated with a diverse group of invertebrate carriers[42, 43]. However, a number of other species in this group have a limited 20 geographic range and form tight associations with specific insect vectors [44, 45]. It is remarkable that the divergent selective regimes associated with these different niches has resulted in such scant morphological change in this group.

*C. elegans* has a famously rigid pattern of development wherein the identity and fate of every cell from the fertilized embryo to the mature adult is known[11]. This set of cell divisions is unchanged across individuals and has allowed the genetic dissection of multiple developmental processes. Yet this developmental system is also highly conserved across multiple genetic backgrounds within species[46-48] and even between species[49]. The highly conserved morphologies in this group are then promoted through highly conserved developmental processes. In tandem with the genetic and ecological diversity observed across the *Caenorhabditis* genus, this is all suggestive of a prevailing role for developmental constraint along its millions of years of evolution.

*C.* sp. 34 clearly bucks this overall pattern, as it displays a morphology and ecology that are distinct departures from its close relatives[12] (Figure 1). Thus developmental constraint alone cannot be driving the patterns of stasis observed in this group. Here, we examined the broad developmental patterns of this divergent species. Together with the extensive background knowledge of the *C. elegans* model system, the roles of constraint and selection in maintaining the general pattern of phenotypic constancy in this group can be interrogated. The existence of mutations in every known protein-coding gene in *C. elegans*[50] provides a window into the universe of evolutionarily-accessible phenotypes that can potentially describe the extent of developmental constraint in this system.

Mutations that affect the body size were among the first described in *C. elegans*[51], and genes that when defective promote long, small, and dumpy (that is, small and fat) phenotypes are among the most notable in this system. Thus, the existence of a novel species that is long (that is, *C.* sp. 34) does not in itself reveal a new region of phenotypic space that was thought to be inaccessible or constrained. However, the developmental biology of these mutants, and their similarity to *C.* sp. 34, reveals insights into the limits of evolutionary trajectories in this group. For instance, given the size difference, it is remarkable that no detectable difference in somatic cell number between *C. elegans* and *C.* sp. 34 was found (Fig. 6A). However, any changes in cell number were also not found when comparing mutants in five body size genes (including both *lon* and *sma*) with wild-type *C. elegans*[52]. This is also consistent with the general observation that nucleus number alone is a poor predictor of body size in rhabditid nematodes[53]. Furthermore, although there are genes that influence cell lineage (and subsequently cell number,[54]), and genes that control germ line proliferation[55], there are, to the best of our knowledge, no mutants with increased body size due to increased cell number in *C. elegans*. Thus, the evolution of body size in *Caenorhabditis* is likely restricted to paths that increase cell size as opposed to cell number. In addition, *C. elegans* body size mutants typically only reveal their differences from wild-type after embryogenesis[56-58]. In *C.* sp. 34, adults are on average 64% longer than *C*. *elegans* adults, but their embryos are only 19% longer (Fig. 5). Thus, both *C. elegans* body size mutants and *C.* sp. 34 largely reveal their differences post-embryonically, which may belie another constraint evolution must operate under to change body size in this group. And further, all of the known long mutants in *C. elegans* do not reveal apparent differences in width[51, 59], the same of which can be said for *C.* sp. 34 and *C. elegans* (Fig. 5k). And finally, to the best of our knowledge, there are no mutants in *C. elegans* that modulate body size by increasing the size of one tissue relative to the others; *C.* sp. 34 likewise reveals a global increase in length (Fig. 6c-d). Thus, when framed within the context of the extensive literature of the *C. elegans* model system, the broad developmental patters of a morphologically divergent close relative can reveal the biases in how certain traits can evolve. But although *C.* sp. 34 appears to be operating under developmental constraints suggested by previous work, its tremendous departure in form from its close relatives remains to be accounted for.

As mentioned above, *Caenorhabditis* species do display diversity in geographic range and phoretic-carrier association. However, it appears that a major aspect of their ecological niche is shared among species in this group: *Caenorhabditis* nematodes generally proliferate on rotting plant material[7, 43]. This likely holds even for extreme invertebrate-vector specialists, although there are reports of necromeny in *C. japonica*[60]. Conversely, *C.* sp. 34 proliferates in the fresh figs of *Ficus septica*[12], the microcosm of which is very different from that of rotting fruit, harboring a unique suite of specific wasps, nematodes, and other microorganisms[61]. This major ecological shift is nearly certain to coincide with a major shift in selective regimes, allowing for the opportunity of novel morphological change in the case of *C.* sp. 34. Thus, the common ecological niches of most *Caenorhabditis* species allow stabilizing selection to maintain a morphology that is suited for rotting-plant bacteriophagy. But, the move to a totally novel niche, as in the case of *C.* sp. 34, has allowed divergent selection to promote novel morphologies within the constraints imposed by its developmental system. Thus selection and constraint act jointly to promote the pattern of morphologies observed in *Caenorhabditis*. In this way, the comparative development approach, together with the context of model systems genetics, can inform long-standing evolutionary questions regarding the interplay of selection and developmental constraint over geological timescales.

## Methods

### Strains

*C.* sp. 34 was originally isolated by NK from a fresh fig of the tree *Ficus septica* in 2013 on Ishigaki Island, Okinawa Prefecture, Japan[12]. The non-isofemale lines NKZ1 and NKZ2 were derived from the same population (also referred to as strain NK74SC), and they are the result of two replicates of an attempt to remove microbial contaminants that have been maintained separately in culture since 2014. Animals were maintained on Nematode Growth Media (with 3.2% agar to discourage burrowing) supplemented with *Escherichia coli* strain OP50-1 for food. *C*. *elegans* strains N2 and JK574 *fog-2 (q71)*[22] were used for most comparisons. Live females/hermaphrodites of *C. briggsae* AF16, *C. remanei* EM464, *C*. *latens* VX88, *C*. *tropicalis* JU1373, *C*. *sinica* JU727, *C. japonica* DF5081 and *C. brenneri* CB5161 were used to illustrate the general morphological constancy of the genus in Figure1

### Size and morphological measurements

For comparing the growth of *C*. *elegans* N2 and *C.* sp. 34 NKZ1 over time (Fig. 5c, animals were synchronized by transferring early-stage embryos to new plates. Each day, a fraction of the synchronized animals was mounted on agar pads in 0.2 mM levamisole, imaged under Nomarski optics, and photographed. Animals were raised at 25°C. L4 *C.* sp. 34 females were moved to a new plate before adulthood in order to prevent mating and the confusion of the synchronized population with their progeny. *C*. *elegans* hermaphrodites were transferred to new plates every day after adulthood for the same purpose. Phenotypically diagnosable males were not used for length measurements. Images were analyzed with the ImageJ software[62] to determine length measurements.

For comparing the sizes of *C*. *elegans* N2 and *C.* sp. 34 NKZ2 at comparable developmental stages (Fig. 5d-k), animals were synchronized by incubating mixed stage animals in a bleaching solution (1 part 10 M KOH: 6 parts sodium hypochlorite: 33 parts water) for seven (*C*. *elegans*) or 4.5 (*C.* sp. 34) minutes. Embryos were then washed four times in M9 buffer and allowed to hatch and arrest in the L1 larval stage overnight at room temperature. Larvae were transferred to bacteria-seeded plates the next day and shifted to 25°C. Observations of developmental timing (described below) were used to determine the timing of the larval stages in *C. elegans* and *C.* sp. 34. Phenotypically diagnosable males were not used for length measurements. Animals at given larval stages were imaged with a dissecting microscope, photographed, and analyzed as above to determine length.

Female/hermaphrodites tail spikes, embryos, and sperm of *C*. *elegans* and *C.* sp. 34 were imaged under Nomarski microscopy and analyzed with ImageJ to quantify morphological differences. Sperm of *C*. *elegans* (*fog-2*) and *C.* sp. 34 NKZ1 males were isolated by cutting off male tails in M9 buffer with a needle.

## Developmental timing

*C. elegans* N2 and *C.* sp. 34 NKZ2 animals were synchronized to the L1 stage as described above. Populations staggered twelve hours apart at 25°C were monitored hourly for the presence of actively molting individuals. Additionally, female vulva and male tail morphology was used to determine the fraction of L4 larvae and adults at a given time. *C*. *elegans* populations were monitored until all individuals developed into adults. *C.* sp. 34 populations were assayed for seven hours after the maximum adult fraction was attained.

## Ploidy, nucleus number, and morphometrics

Ploidy and nucleus observations were made using animals stained with the DNA-staining Hoechst 33342 dye. *C. elegans fog-2 (q71)* and *C.* sp. 34 NKZ2 young adult females were obtained by moving L4 females to new plates at 25°C and preparing them one (*C*. *elegans*) or two (*C.* sp. 34) days later for fluorescence microscopy. Animals were then fixed in 100% methanol for ten minutes at-20°C. Animals were washed three times in PBS and were then incubated in 1 µg/ml Hoechst 33342 for 10 minutes. Animals were washed three times in PBS and then mounted in 50% glycerol for visualization. Specimens for cell number, ploidy, and morphometrics were imaged with an Olympus FluoView 1000 laser-scanning confocal microscope. A fraction of specimens were examined for ploidy using a conventional compound microscope equipped with fluorescence.

For the determination of ploidy, proximal oocytes in prophase I arrest were imaged and diakinesis chromosomes counted. For the determination of somatic nucleus number, z-stacks with one micron steps across the whole specimen were generated. All somatic nuclei were then hand counted using the cell counter plugin in the ImageJ software. ImageJ was then used to also determine the distance between homologous morphological landmarks using the same sets of images. Homologous landmarks were determined by anatomical similarity and the relative positions of nuclei. The morphological landmarks used were: the most anterior nucleus observed (likely Hyp4, number 1 on Figure 6b); the most anterior intestinal nucleus (Int4, #2); the BDUL neuron (#3); the anterior spermatheca (measured at the end of the proximal-1 oocyte, #4); the center of the vulva (#5); the posterior spermatheca (measured at the end of the proximal-1 oocyte, #6); the VD11 neuron (#7); the most posterior intestinal nucleus (Int9, #8); and the most posterior nucleus (likely Hyp10, #9).

## Acknowledgements

Some strains were provided by the CGC, which is funded by National Institutes of Health Office of Research Infrastructure Programs (P40 OD010440). We thank N. Kanzaki for discovering and sharing *C.* sp. 34. We thank K. Kuroda, B. Armstrong, K. Prehoda, and D. Libuda for assistance in fluorescence and confocal microscopy. We thank T. Kikuchi for sharing the *C.* sp. 34 genome. We thank E. Schwarz for sharing the *C. wallacei* genome. Funding for this work was provided by the Japan Society for the Promotion of Science and the National Institutes of Health (5F32GM115209-03; R01GM102511).

## Author Contributions

GCW conducted and analyzed the experiments; GCW and PCP designed the experiments; GCW and PCP wrote the paper.

## Competing Financial Interests

The authors declare no competing financial interests.

## Materials and Correspondence

Correspondence and requests for materials should be addressed to Gavin Woodruff at.

---

Supplemental Figure 1. Germ line abnormalities in *C*. sp. 34.

Supplemental Table 1. FASconCAT supermatrix information.

Supplemental Table 2. Descriptions of protein sequences used for phylogenetic analysis.

Supplemental Document 1. Discussion of a phylogenetic analysis.

Supplemental Document 2. Description and results of interspecies mating tests.

Supermatrix.phy. Alignment used for phylogenetic analysis.

Supermatrix_partition.txt. Partition file for phylogenetic analysis.

Raxml_input.txt. Command-line input for RAxML.

